# WFIKKN2 is a bifunctional axon guidance cue that signals through divergent DCC family receptors

**DOI:** 10.1101/2023.06.15.544950

**Authors:** Kelsey R. Nickerson, Irene Tom, Elena Cortés, Jane R. Abolafia, Engin Özkan, Lino C. Gonzalez, Alexander Jaworski

## Abstract

Axon pathfinding is controlled by attractive and repulsive molecular cues that activate receptors on the axonal growth cone, but the full repertoire of axon guidance molecules remains unknown. The vertebrate DCC receptor family contains the two closely related members DCC and Neogenin with prominent roles in axon guidance and three additional, divergent members – Punc, Nope, and Protogenin – for which functions in neural circuit formation have remained elusive. We identified a secreted Punc/Nope/Protogenin ligand, WFIKKN2, which guides mouse peripheral sensory axons through Nope-mediated repulsion. In contrast, WFIKKN2 attracts motor axons, but not via Nope. These findings identify WFIKKN2 as a bifunctional axon guidance cue that acts through divergent DCC family members, revealing a remarkable diversity of ligand interactions for this receptor family in nervous system wiring.

**One-Sentence Summary:** WFIKKN2 is a ligand for the DCC family receptors Punc, Nope, and Prtg that repels sensory axons and attracts motor axons.

## Main Text

Neural circuit assembly during embryonic development requires the guidance of extending axons to their proper targets. Axon pathfinding is orchestrated by attractive and repulsive molecular cues that are sensed by receptors on the leading process of the axon, the growth cone *(1)*. While numerous ligand-receptor pairs have been implicated in steering axon growth, the full repertoire of axon guidance molecules likely includes additional, yet-to-be-identified cues and receptors.

The receptor deleted in colorectal cancer (DCC) and its ligand Netrin-1 mediate axon attraction and instruct neuronal wiring in various organisms, including worms, flies, mice, and humans *(2–6)*. DCC is a type I transmembrane protein with an N-terminal extracellular domain (ECD) that contains four immunoglobulin-like (IG) domains followed by six fibronectin type III (FN) domains; the fourth through sixth FN domains interact with the N-terminal laminin (LN) and the three epidermal growth factor-like (EGF) domains of Netrins (Fig. 1A) *(4, 7–9)*. In the DCC intracellular domain (ICD), three conserved motifs, P1-P3, have been implicated in mediating Netrin signaling via recruitment of various kinases *(10–15)*. In vertebrates, a structurally similar member of the DCC family, Neogenin (Neo1) (Fig. 1A), also mediates attraction to Netrin *(4, 8, 16)*. DCC and Neo1 further interact with other ligands, which they do not share, e.g., DCC binds the guidance cue Draxin *(17)*, and Neo1 binds repulsive guidance molecule (RGM) family proteins *(18, 19)*. The vertebrate DCC family contains three additional receptors (Fig. 1A) *(20)*: Punc (also known as Igdcc3) *(21)*, Nope (Igdcc4) *(22)*, and Protogenin (Prtg, Igdcc5) *(23)*. These three receptors differ from DCC and Neo1 in their number of FN domains, with Prtg and Nope having five and Punc having two, and their ICDs lack the P motifs found in DCC/Neo1. While expression of Punc, Nope, and Prtg has been reported in the mouse and chick embryonic nervous system *(21–25)*, no functional ligands for these receptors have been identified, and possible roles for these orphan DCC family members in axon guidance and neural circuit formation have remained elusive.

**Fig 1.**
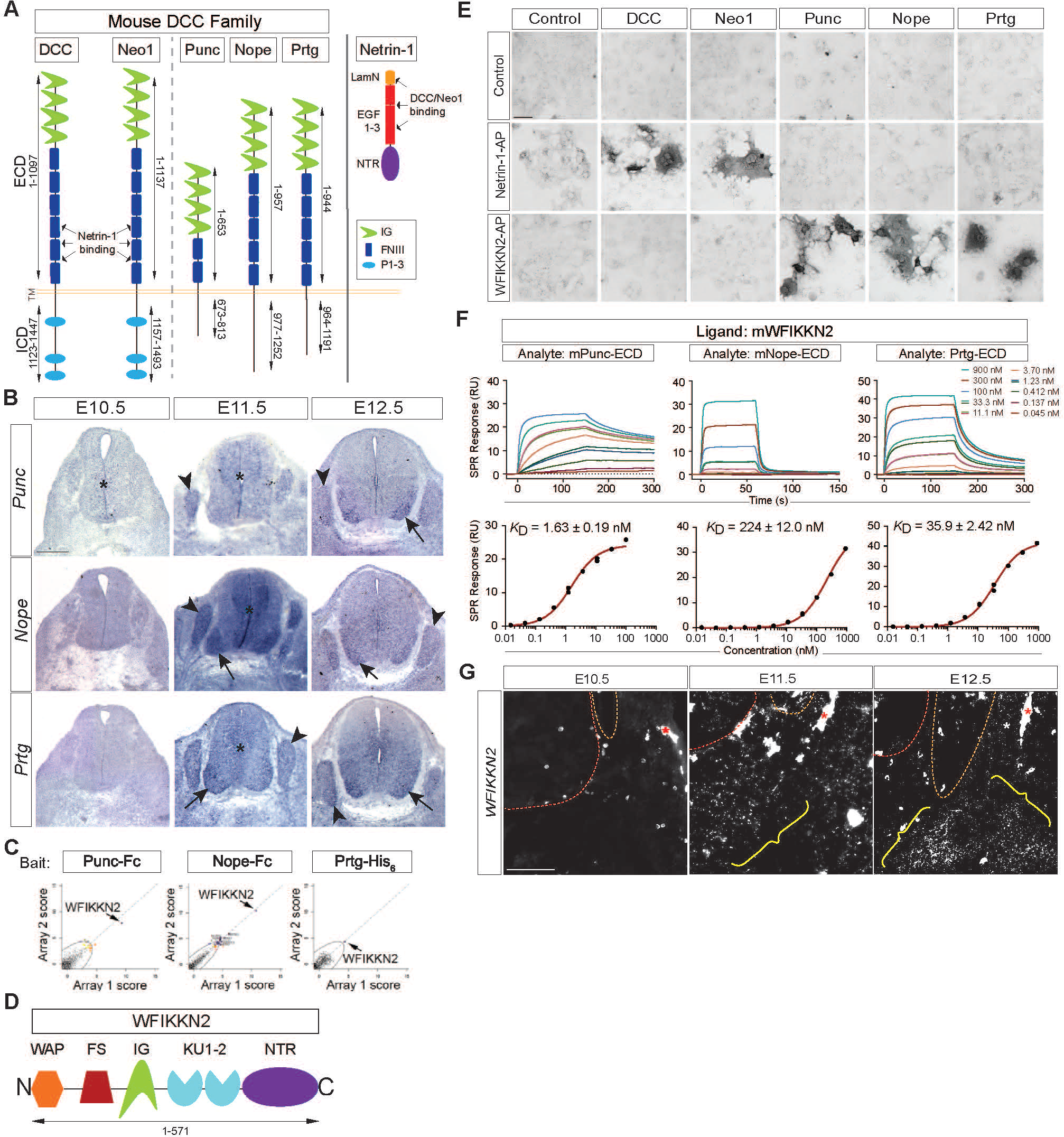
Punc, Nope, and Prtg are expressed in developing sensory and motor neurons and bind WFIKKN2, expressed in the spinal cord periphery. **(A)** Domain structure of murine (m)DCC family receptors and Netrin-1, drawn to scale. TM, transmembrane domain. **(B)** Brachial spinal cord transverse sections from E10.5, E11.5, and E12.5 mouse embryos were used for chromogenic in situ hybridization to detect *Punc, Nope,* and *Prtg* mRNA. All three genes show strong expression in DRGs (arrowheads) and the ventral horn (arrows) beginning at E11.5, with expression of *Punc* in ventral horn slightly delayed. *Punc* is also expressed in the progenitor zone (asterisk) at E10.5; all three receptors are expressed in the progenitor zone at E11.5 **(C)** An extracellular protein microarray was screened using Punc-ECD-Fc, Nope-ECD-Fc, and Prtg-ECD-His_6_ as baits, revealing that all three receptors bind WFIKKN2. Black dots represent results for individual prey proteins from duplicate screening (arrays 1, 2). **(D)** Domain structure of mWFIKKN2. **(E)** Interactions of DCC family receptors with Netrin-1 and WFIKKN2 were examined in a COS-7-based binding assay. Netrin-1-AP specifically binds cells expressing DCC or Neo1, but not cells expressing Punc, Nope, or Prtg. Conversely, WFIKKN2-AP binds Punc-, Nope-, or Prtg-expressing cells, but not cells expressing DCC or Neo1. **(F)** WFIKKN2-Punc/Nope/Prtg binding was studied by SPR. Shown are SPR sensograms and Langmuir isotherms of steady-state responses fitted to a specific binding model for mWFIKKN2 as ligand on sensor chips with mPunc, mNope, or mPrtg ECDs as analytes at indicated concentrations. *K*_D_ values are indicated, including standard error of the model fit. **(G)** Brachial-level transverse sections of E10.5, E11.5, and E12.5 embryos were used for RNAScope fluorescent in situ hybridization to detect *WFIKKN2* mRNA. *WFIKKN2* is expressed in tissue (yellow brackets) surrounding the spinal cord and DRG (red and orange dotted outlines, respectively) from E11.5 onwards. Asterisks indicate non-specific labeling from blood vessels. Scale bars, (B) 200 μm; (E) 50 μm; (G) 100 μm. Error bars indicate SEM.

### *Punc*, *Nope*, and *Prtg* are expressed in developing sensory and motor neurons

Previous studies reported expression of Punc, Nope, and Prtg in the developing spinal cord and dorsal root ganglia (DRGs) *(21–24, 26)*. We therefore examined *Punc, Nope,* and *Prtg* expression in the spinal cord and DRG (fig. S1A) during the time of DRG sensory axon growth into the body periphery [embryonic day (E)10.5 to 14.5] *(27)* by means of in situ hybridization (ISH). At E10.5, *Punc* is expressed in the ventricular zone of the spinal cord, while *Nope* and *Prtg* are undetectable (Fig. 1B). Beginning at E11.5 and persisting through E14.5, all three receptor genes are expressed in DRGs (Fig. 1B, fig. S1B), coinciding with the extension of central and peripheral sensory axon branches *(28)*. The receptors are also expressed in the spinal cord ventral horn, where motor neurons reside (fig. S1A). Here, *Nope* and *Prtg* expression begins at E11.5, after motor axons have exited the spinal cord and are en route to their targets, and persists through E14.5, when many motor axons have reached their target muscles; *Punc* mRNA is also detectable in the ventral horn, but only from E12.5 onwards (Fig. 1B, fig. S1B) *(30)*. To validate expression of Punc, Nope, and Prtg in motor and DRG sensory neurons, we analyzed existing E12.5 single-cell RNA sequencing (scRNA-seq) data *(29)* for the presence of receptor mRNAs in these cell types. We confirmed the presence of *Punc, Nope,* and *Prtg* transcripts, either individually or in various combinations, in motor and sensory neurons; this includes motor neurons belonging to the medial and lateral motor column (MMC and LMC, respectively) and various sensory neuron subtypes defined by tropomyosin receptor kinase A (TrkA), TrkB, and/or TrkC expression (fig. S1C, D) *(29–31)*. Thus, *Punc*, *Nope*, and *Prtg* are expressed in spinal motor neurons and DRG sensory neurons during key developmental timepoints of axon extension, suggesting that these receptors may control neuronal wiring.

### WFIKKN2 binds Punc, Nope, and Prtg and is expressed in the spinal cord periphery

To identify ligands that might engage Punc, Nope, and Prtg for axon guidance, we screened secreted protein microarray *(32)* with each of the receptors’ ECDs fused to Fc or a hexahistidine tag (Punc-ECD-Fc, Nope-ECD-Fc, Prtg-ECD-His_6_) and found that all three receptors bind a fusion of WFIKKN2 and Fc (Fig. 1C). WFIKKN2 (also known as GASP-1) is a secreted protein containing a whey acidic protein (WAP) domain, a follistatin (FS) domain, an IG domain, two Kunitz-type protease inhibitor (KU) domains, as well as a Netrin (NTR) module (Fig. 1D) *(33, 34)*. WFIKKN2 can bind and inhibit multiple members of the transforming growth factor-β (TGFβ) family to regulate muscle growth and skeletal patterning *(35–38)*, but functions for WFIKKN2 in neural circuit formation have not been reported. To confirm the results of the microarray screen and delineate the binding specificities of DCC family members for Netrin-1 and WFIKKN2, we transfected COS-7 cells with expression constructs for DCC, Neo1, Punc, Nope, or Prtg and tested for binding of alkaline phosphatase (AP) fusions of Netrin-1 or WFIKKN2 (Netrin-1-AP, WFIKKN2-AP). We found that Netrin-1-AP binds cells expressing DCC or Neo1, but not the other three receptors, while WFIKKN2-AP binds Punc, Nope, and Prtg, but not DCC or Neo1 (Fig. 1E). We confirmed the interactions between WFIKKN2 and Punc/Nope/Prtg by means of surface plasmon resonance (SPR), which yielded *K*_D_ values in the 1.63-224 nM range (Fig. 1F), while binding between WFIKKN2 and DCC/Neo1 is undetectable (fig. S1E). Thus, Punc, Nope, and Prtg bind WFIKKN2 with high affinity, and Netrin-1 and WFIKKN2 interact with non-overlapping subsets of DCC family receptors.

To determine whether the axons of motor and sensory neurons, which express Punc, Nope, and Prtg, encounter WFIKKN2 in vivo, we used fluorescent ISH and examined *WFIKKN2* expression in the spinal cord and surrounding tissues during the time when these axons navigate through the periphery [E10.5-E12.5]. At E10.5, *WFIKKN2* mRNA is undetectable in the neural tube and its vicinity (Fig. 1G), but at E11.5 and E12.5, *WFIKKN2* is expressed in the presumptive sclerotome bordering the ventral spinal cord where motor and sensory axons coalesce at the motor-sensory plexus (MSP) and subsequently form the dorsal and ventral ramus (Fig. 1G, fig. S1A, S1F) *(39)*. *WFIKKN2* mRNA is excluded from the spinal cord and DRG during this time and undetectable in *WFIKKN2* knockout embryos (Fig. 1G, fig. S1F), demonstrating specificity of the ISH results. Hence, the axons of motor neurons and DRG sensory neurons, which express WFIKKN2 receptors, extend through areas of WFIKKN2 expression in the spinal cord periphery.

### WFIKKN2 repels DRG sensory axons and contributes to their guidance in vivo

The interaction between WFIKKN2 and Punc/Nope/Prtg and the complementary expression patterns of this ligand and its receptors suggest that WFIKKN2 might regulate motor and sensory axon guidance in the spinal cord periphery. Because the timing of *WFIKKN2* expression adjacent to the neural tube aligns more closely with the growth of sensory axons, which follow earlier-extending motor axons into the body periphery *(39)*, we first examined the response of sensory axons to WFIKKN2 in vitro. DRG explants from E12.5 mouse embryos were cultured with nerve growth factor (NGF) to promote axon extension from sensory neurons expressing the NGF receptor TrkA *(40)*, which constitute the majority of DRG neurons at this stage of development *(30)*. When confronted with aggregates of COS-7 cells that coexpress red fluorescent protein (RFP) and WFIKKN2-AP, sensory axons grow away from the cell aggregates, whereas axon outgrowth from DRG explants remains radially symmetric in the presence of RFP-expressing (control) cells (Fig. 2A, B). This repulsive effect of WFIKKN2 was confirmed when we adapted Dunn chamber axon turning assays *(41)* to study responses of individual E11.5 +1 day in vitro (DIV) TrkA-expressing DRG neurons to gradients of WFIKKN2; here, WFIKKN2 causes axon repulsion at peak concentrations of 100 ng/ml and above (Fig. 2C, D, movies S1-S2). Thus, as shown in two different guidance assays, WFIKKN2 repels sensory axons in vitro.

**Fig 2.**
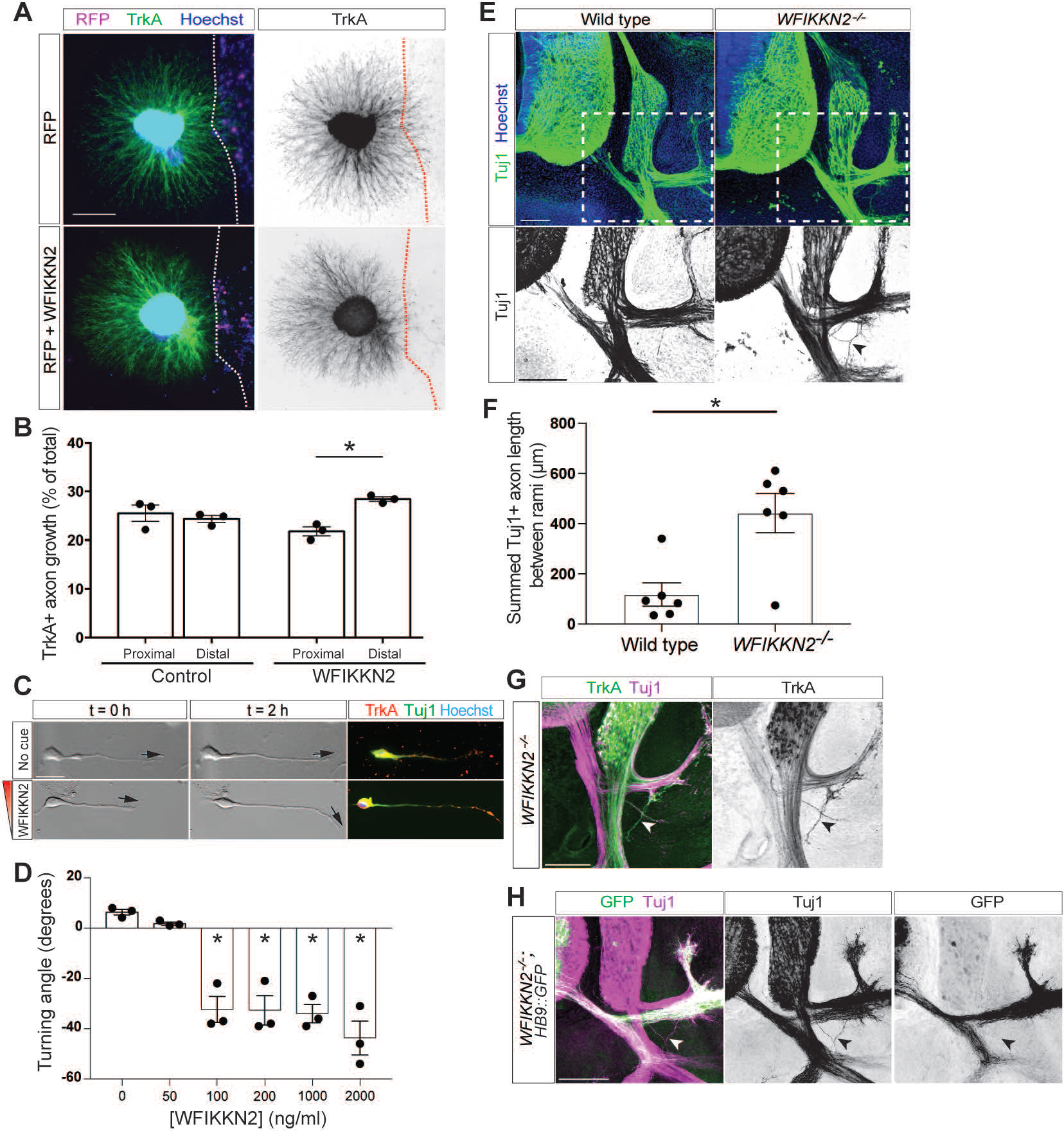
WFIKKN2 repels sensory axons and prevents them from leaving the main axon bundles in the spinal cord periphery. (**A**) E12.5 mouse DRG explants were cocultured with COS-7 cells (dotted outline) that express RFP or RFP and WFIKKN2-AP and stained for TrkA to visualize sensory axons. Hoechst stain labels cell nuclei. WFIKKN2 expression causes axon turning away from cell aggregates. (**B**) Sensory axon growth in the proximal and distal quadrants relative to COS-7 aggregates was quantified. With WFIKKN2-expressing cells, distal growth is significantly higher than proximal growth (n = 3 independent experiments, *p* = 0.0409), while proximal and distal growth are comparable in the presence of control cells (n = 3, *p* = 0.3681). (**C**) Dissociated E11.5 + 1 DIV DRG neurons were exposed to a WFIKKN2 gradient or no guidance cue in Dunn chambers. DIC images at 0 (left) and 2 hours (middle) show sensory axon turning away from the WFIKKN2 gradient. Black arrows indicate orientation of distal axon tip. Post-hoc immunohistochemistry (IHC) for TrkA and Tuj1 (right) confirms molecular identity of analyzed neurons. (**D**) Quantification of sensory axon turning angles for increasing concentrations of WFIKKN2 (n = 3 independent experiments for all conditions). Negative turning angles for concentrations ≥ 100 ng/ml indicate axon repulsion (comparison to control: 100 ng/ml *p* = 0.0006, 200 ng/ml *p* = 0.0006, 1000 ng/ml *p* = 0.0005, 2000 ng/ml *p* < 0.0001). (**E**) Brachial-level transverse sections of E12.5 wild-type and *WFIKKN2^−/−^* embryos were labeled with Tuj1 antibody and Hoechst stain. In both genotypes, axons are organized into the dorsal and ventral ramus distal to the MSP in the spinal cord periphery; in *WFIKKN2*^−/−^ but not wild-type mice, a subset of axons defasciculates from the main nerve bundles and projects into the area between the two rami (black arrowhead). (**F**) Ectopic axon growth at the MSP was quantified and is significantly higher in *WFIKKN2^−/−^* embryos thank in wild type (n = 6 embryos/genotype, *p* = 0.0052). (**G**) E12.5 *WFIKKN2^−/−^*transverse embryo sections were stained for TrkA and Tuj1. Misprojecting axons (black arrowhead) at the MSP are TrkA-positive sensory axons. (**H**) E12.5 *WFIKKN2^−/−^;HB9::GFP* embryo sections were stained for Tuj1 and GFP. GFP-positive motor axons do not stray from the main nerve bundles at the MSP. Scale bars, (A, E, G, H,) 100 μm;(C) 25 μm. Error bars indicate SEM.

To determine whether WFIKKN2 controls axon pathfinding in vivo, we analyzed mice deficient in *WFIKKN2 (42)*. We stained transverse spinal cord sections from E10.5, E11.5, and E12.5 *WFIKKN2^−/−^* embryos and their wild-type littermates with antibodies against the panaxonal marker class III β-tubulin (Tuj1) and examined the formation of axon tracts in the spinal cord periphery. In E10.5 wild-type embryos, efferent motor axons and the peripheral branches of DRG sensory axons have formed the MSP, and a subset of sensory axons and LMC motor axons project into the developing ventral ramus, while a small number of MMC motor axons begin to extend epaxially (fig. S1A, S2A) *(39)*; at E11.5, the dorsal ramus has formed (fig. S1A, S2A), and growth of additional motor and sensory axons between E11.5 and E12.5 leads to expansion of the ventral and dorsal ramus (Fig. 2E, fig. S1A, S2A) *(39).* In *WFIKKN2^−/−^* embryos, peripheral projections appear normal at E10.5 and E11.5 (fig. S2A); however, at E12.5, a subset of axons has defasciculated from the main nerve branches at the MSP and projects into the area between the ventral and dorsal ramus, often in a web-like pattern (Fig. 2E, F). Antibody labeling for the neurotrophin receptors TrkA, TrkB, and TrkC, which mark different subsets of DRG sensory neurons *(43)*, revealed that misprojecting axon bundles in *WFIKKN2^−/−^*embryos contain TrkA- and TrkC-positive, but not TrkB-positive axons (Fig. 2G, fig. S2B, C); this phenotype is detectable at all rostro-caudal levels (data not shown). We also examined *WFIKKN2^−/−^*embryos carrying the motor neuron-specific *HB9::GFP* transgene and found that motor axons do not join the misguided sensory axons and instead remain within the ventral and dorsal ramus (Fig. 2H). Thus, WFIKKN2 is required to prevent defasciculation and ectopic projection of sensory axons at the MSP. Consistent with lack of detectable *WFIKKN2* expression in DRGs and further arguing against a requirement for sensory neuron-derived WFIKKN2 in fasciculation, sensory axon growth and bundling from *WFIKKN2^−/−^* DRG explants cultured in vitro is indistinguishable from wild-type explants (fig. S2D, E). Together, these results support the idea that repulsion from peripherally expressed WFIKKN2 forces sensory axons to remain within the main nerve bundles at the MSP.

### Nope mediates sensory axon guidance by WFIKKN2

We next tested whether WFIKKN2 repulsion of sensory axons is mediated by Punc/Nope/Prtg. When we performed Dunn chamber axon turning assays with DRG sensory neurons in gradients of WFIKKN2 and bath-applied Nope-ECD-Fc as a competitor for WFIKKN2 binding to its endogenous receptors, we found that WFIKKN2-mediated repulsion is completely blocked (Fig. 3A, B). This result supports the idea that WFIKKN2 effects sensory axon repulsion via a receptor that binds the same site as Nope-ECD-Fc, including Punc, Prtg, and Nope itself.

**Fig. 3.**
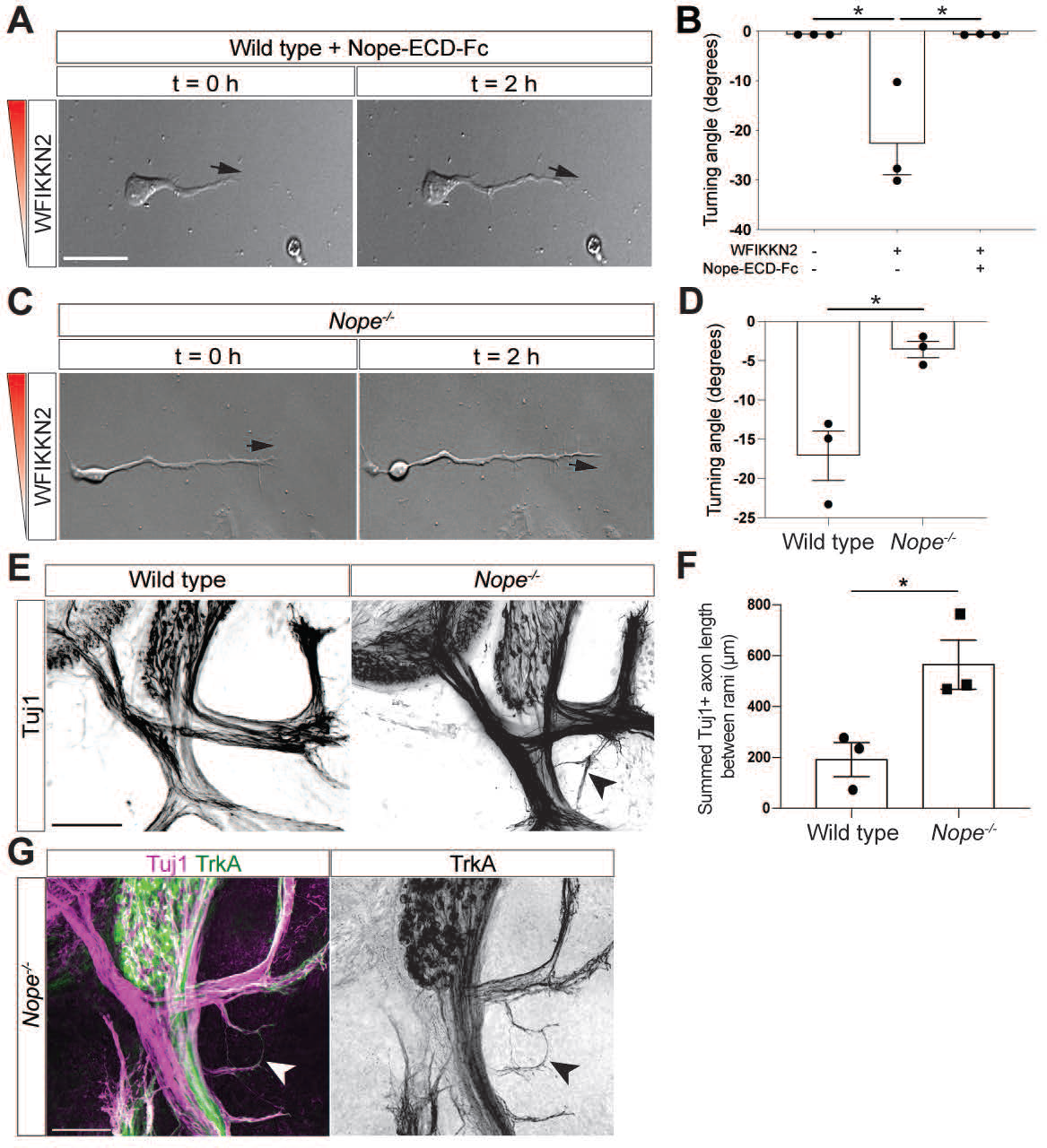
WFIKKN2 guides sensory axons through Nope. (**A**) Dunn chamber axon turning assays using E11.5 wild-type DRG neurons. Incubation with 25 μg/ml Nope-ECD-Fc prevents sensory axon turning in response to 100 ng/ml WFIKKN2. Black arrows indicate orientation of distal axon tip. (**B**) Quantification of sensory axon turning angles shows complete block of repulsion from WFIKKN2 by Nope-ECD-Fc bath application (n = 3 experimental replicates, no cue vs. WFIKKN2 *p =* 0.0119, WFIKKN2 vs. WFIKKN2 + Nope-ECD-Fc *p* = 0.0119). (**C**) Dunn chamber axon turning assays using E11.5 *Nope^−/−^* DRG neurons. Genetic inactivation of *Nope* prevents sensory axon turning in response to 100 ng/ml WFIKKN2. Black arrows indicate orientation of distal axon tip. (**D**) Quantification of *Nope*^−/−^ and wild-type littermate sensory axon turning angles shows that repulsion from WFIKKN2 is abolished in *Nope*^−/−^ neurons (n = 3 biological replicates, *p =* 0.0152). (**E**) Brachial-level transverse sections of E12.5 *Nope^−/−^* embryos and their wild-type littermates were labeled with Tuj1 antibody and Hoechst stain. Stray axons (black arrowhead) invade the area between the dorsal and ventral ramus in *Nope^−/−^* mice but not wild type. (**F**) Quantification reveals significantly increased axon growth between the two rami in *Nope^−/−^* embryos compared to wild type (n = 3 embryos per genotype, *p* = 0.03326). (**G**) E12.5 *Nope^−/−^* embryo sections were stained for TrkA and Tuj. Misprojecting axons (arrowhead) are TrkA-positive sensory axons. Scale bars, (A) 25 μm; (C,E) 100 μm. Error bars indicate SEM.

We then used genetic approaches to probe the roles of individual DCC family receptors in WFIKKN2-mediated repulsion. We first generated mice deficient in *Nope* (fig. S3A) and confirmed via quantitative reverse transcription polymerase chain reaction (qRT-PCR) that *Nope* mRNA is absent or dramatically (96-99%) reduced in these animals (fig. S3B), indicating that the mutant allele is null. When we exposed DRG neurons from *Nope^−/−^* and wild-type littermate embryos to WFIKKN2 in Dunn chambers, we observed robust repulsion of wild-type axons but found that *Nope* mutant sensory neurons fail to respond to WFIKKN2 (Fig. 3C, D). Thus, Nope is required for sensory axon repulsion by WFIKKN2 in vitro.

To determine whether Nope, like WFIKKN2, is required for sensory axon guidance in vivo, we analyzed peripheral axon projections in *Nope^−/−^* embryos by immunohistochemistry (IHC). We observed that TrkA- and TrkC-positive, but not TrkB-positive sensory axons defasciculate from the main nerve bundles at the MSP in these mice and ectopically project into the region between the dorsal and ventral ramus (Fig. 3E-G, fig. S3C, D). The severity of this defect is indistinguishable from the *WFIKKN2^−/−^*phenotype (compare Figs. 2F and 3F). Thus, Nope aids in sensory axon bundling at the MSP, supporting the idea that it mediates the function of WFIKKN2 in guiding these axons through repulsion.

### WFIKKN2 attracts motor axons

Guidance of DRG sensory axons through the body periphery is strongly influenced by interactions with earlier-extending motor axons, which provide a scaffold for sensory axon growth *(39)*. Most motor axons emerge from the neural tube before the onset of *WFIKKN2* expression in the spinal cord periphery at E11.5 (*27, 39)*, and misprojecting sensory axons in mice lacking WFIKKN2 are not accompanied by motor axons (Fig. 2H). This argues that sensory axon guidance defects in mice with disrupted WFIKKN2-Nope signaling are not a secondary consequence of earlier mistakes made by motor axons. Importantly, is also raises the interesting question why late-extending motor axons, which express Punc/Nope/Prtg and encounter WFIKKN2, do not stray from their path in *WFIKKN2^−/−^* embryos, similar to sensory axons. To address this question, we tested whether WFIKKN2 acts as a repellant for motor axons in vitro. Ventral spinal cord explants from E11.5 mouse embryos were cultured under conditions that promote radially symmetric outgrowth of motor axons *(44)*. When cocultured with cells that express either WFIKKN2-AP and RFP or RFP alone, motor axons preferentially grow towards WFIKKN2-expressing cells, but not control cells (Fig. 4A, B), while overall axon outgrowth is not significantly increased by WFIKKN2 (normalized total axon length, 100 ± 27.8% for control, 84 ± 19.7% for WFIKKN2; *p* = 0.3436, n = 3 experiments). We confirmed this attractive effect in Dunn chamber axon turning assays with dissociated E10.5 +1 DIV ventral horn neurons, where WFIKKN2 elicits positive turning responses from axons of Islet1/2-expressing motor neurons at peak concentrations of 500 ng/ml and above (Fig. 4C, D, movies S3, S4). Axon attraction to WFIKKN2 in Dunn chambers was observed for both MMC and LMC neurons (fig S4). Thus, WFIKKN2 attracts rather than repels motor axons in vitro.

**Fig 4.**
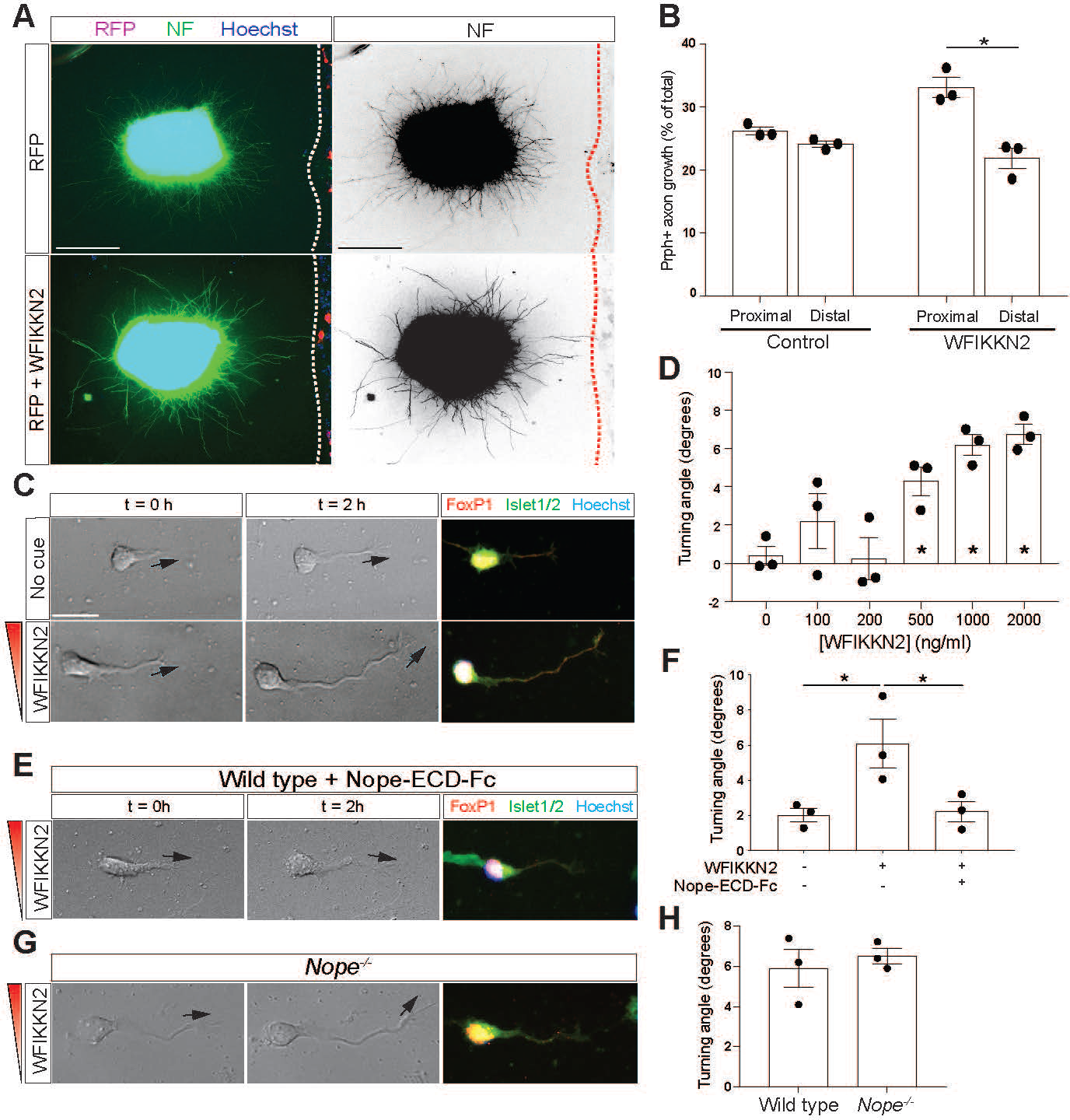
WFIKKN2 attracts motor axons in vitro, independent of Nope. (**A**) E11.5 mouse ventral horn explants were cocultured with COS-7 cells (dotted outline) that express RFP or RFP and WFIKKN2 and labeled with neurofilament (NF) antibody to visualize axons, as well as Hoechst stain. (**B**) Axon outgrowth in the proximal and distal quadrants relative to COS-7 aggregates was quantified. With WFIKKN2-expressing cells, proximal growth is significantly higher than distal growth (n = 3*, p* = 0.0149), while proximal and distal growth are comparable in the presence of control cells (n = 3, *p* = 0.5593). (**C**) Dissociated E10.5 + 1 DIV ventral spinal cord neurons were exposed to a WFIKKN2 gradient (1000 ng/ml) or no guidance cue in Dunn chambers. DIC images at 0 and 2 h show motor axon turning towards the WFIKKN2 gradient. Black arrows indicate orientation of distal axon tip. Post-hoc IHC for FoxP1 and Islet1/2 (right) confirms molecular identity of analyzed neurons. (**D**) Quantification of motor axon turning angles for increasing concentrations of WFIKKN2 (n = 3 independent experiments for all conditions). Positive turning angles for concentrations ≥ 500 ng/ml indicate axon attraction (comparison to control: 500 ng/ml *p* = 0.0321, 1000 ng/ml *p* = 0.0080, 2000 ng/ml *p* = 0.0053). (**E)** Dunn chamber axon turning assays using E10.5 wild-type motor neurons. Incubation with 25 μg/ml Nope-ECD-Fc prevents motor axon turning in response to 1000 ng/ml WFIKKN2. Black arrows indicate orientation of distal axon tip. (**F**) Quantification of motor axon turning angles shows inhibition of attraction to WFIKKN2 by Nope-ECD-Fc application (n = 3, no cue vs. WFIKKN2 *p =* 0.0361, WFIKKN2 vs. WFIKKN2 + Nope-ECD-Fc *p* = 0.0356). (**G**) Dunn chamber axon turning assays using E10.5 *Nope^−/−^* motor neurons. Genetic inactivation of *Nope* does not eliminate motor axon turning response to 1000 ng/ml WFIKKN2. Black arrows indicate distal axon tip. (**H**) Quantification of *Nope^−/−^* and wild-type littermate motor axon turning angles shows comparable attraction to WFIKKN2 in the two genotypes (n = 3, *p =* 0.5860). Scale bars, (A) 100 μm; (C, E) 25 μm. Error bars indicate SEM.

To determine how motor axon attraction to WFIKKN2 is mediated, we again used Nope-ECD-Fc as a function-blocking reagent in Dunn chamber axon turning assays and found that it abolishes motor axon responses to WFIKKN2 (Fig. 4E, F), consistent with the idea that WFIKKN2 attracts motor axons through Punc, Nope, or Prtg. We also studied the axon turning responses of motor neurons from *Nope* mutant animals and found that motor axons are still attracted to WFIKKN2 (Fig. 4G, H). Taken together, these data indicate that WFIKKN2’s attractive effect on motor axons does not require Nope and is instead likely mediated by Punc or Prtg.

## Discussion

Our results identify WFIKKN2 as a bifunctional guidance cue that repels sensory axons and attracts motor axons. WFIKKN2 binds the three divergent DCC family receptors Punc, Nope, and Prtg, and WFIKKN2-Nope repulsive signaling contributes to the guidance of sensory axons in the spinal cord periphery by preventing them from leaving the main nerve bundles. This type of repellant-mediated mechanism for forcing axon fasciculation is commonly termed “surround repulsion” *(45, 46)*. Motor axon attraction to WFIKKN2 can be blocked by competing for binding to Punc/Nope/Prtg, but not by genetic inactivation of Nope, implying that Punc or Prtg are mediators of WFIKKN2 attraction. This suggests a model in which WFIKKN2 engages different, yet related, receptors to elicit either axon attraction or repulsion in specific neurons, possibly via recruitment of distinct coreceptors or downstream signaling molecules. While a recent cell-surface interactome screen implicated WFIKKN2 as a binding partner for all five DCC family receptors *(20, 47)*, our results, from two independent types of binding assays, demonstrate that Netrin-1 receptors [DCC and Neo1] and WFIKKN2 receptors [Punc, Nope, and Prtg] form two mutually exclusive subgroups within the vertebrate DCC family. While both ligands instruct axonal pathfinding, this reveals a striking functional specialization among the receptors, which is likely to contribute to the increase in nervous system complexity that accompanied evolution of the DCC family.

## Acknowledgments

We thank Y. Zhou for expert technical assistance and the Brown University Mouse Transgenic and Gene Targeting Facility for assistance with generation of the *Nope* mutant mice. We are grateful to S.-J. Lee for sharing the *WFIKKN2/Gasp1* mutant mouse line and T. Suter for providing probe templates for *Punc/Nope/Prtg* in situ hybridization, and we are also thankful to G. Barnea and members of the Jaworski and Özkan laboratories for thoughtful comments on the manuscript.

## Author contributions

Conceptualization: AJ

Investigation: KN, IT, EC, JA

Supervision: EÖ, LG, AJ

Visualization: KN, IT, EC, JA

Formal Analysis: KN, JA, EC, IT

Writing – original draft: KN, AJ

Writing – review & editing: KN, IT, EC, JA, EÖ, LG, AJ

## Data and materials availability

All data are available in the main text or the supplementary materials.

## Supplementary Materials

Materials and Methods

Figs. S1 to S4

Table S1

References (*48–53*)

Movies S1 to S4

## Materials and Methods

### Animals

All experimental procedures had institutional approval through Brown University’s Institutional Animal Care and Use Committee and followed the guidelines provided by the National Institutes of Health. Mice carrying the *WFIKKN2* null allele (MGI:5546173) *(42)* and *HB9::GFP* transgene (MGI: 3056906, official nomenclature Tg(Hlxb9-GFP)1Tmj) *(48)* have been described before and were genotyped by PCR as originally reported. Mice carrying the *Nope^−^*null allele were generated through targeted insertion of an in-frame stop codon (TAA) followed by an IRES-mCherry-poly(A) expression cassette into exon 2 of the *Nope* locus by homology-directed repair after introduction of a double strand break via CRISPR/Cas9. The *Nope^+^* and *Nope^−^* alleles were detected by PCR from genomic DNA (primer sequences: GAAGTAAGGGGCTGTTGGCA, ACACCGAGCAGCAAGAGTAG, and GTGCCGCCTTTGCAGGTGTATC). Mice were maintained on a CD-1 background (wild-type, *WFIKKN2* mutant, and *HB9::GFP* transgenic lines) or a mixed CD-1/C57BL6 background (*Nope* knockout line). For timed pregnancies, the day of vaginal plug was defined as E0.5, and littermate embryos of either sex were used for all experiments.

### In situ hybridization

Chromogenic in situ hybridization on 20-μm-thick transverse embryo sections (brachial level) was essentially carried out as described previously *(49)*, with a hybridization temperature of 65°C and post-hybridization wash temperature of 55 °C. The used riboprobes span nucleotides 314 through 1238 of the *Punc* coding sequence (cds) (GenBank accession NM_001357257.1), nucleotides 487 through 1148 of the *Nope* cds (NM_001290315), and nucleotides 417 through 1385 of the *Prtg* cds (NM_175485). Labeling with sense probes resulted in weak, uniform staining for each gene (fig. S1B). For RNAscope fluorescent in situ hybridization, slides with 20-μm-thick transverse embryo sections (brachial level) were processed according to the ACD RNAscope fresh frozen tissue pretreatment and fluorescent multiplex assay manual. The Mm-WFIKKN2 probe (generated by ACD Bio) was used in accordance with the manufacturer’s recommendations. RNAscope was combined with immunohistochemistry (IHC) as previously described *(11). WFIKKN2* in situ hybridization on spinal cord sections from E12.5 *WFIKKN2^−/−^* embryos did not produce detectable signal (fig. S1F). Images were acquired on a Nikon Ti-E microscope.

### Analysis of published scRNA-Seq data

A published single-cell transcriptomic dataset of embryonic spinal cord and surrounding tissues *(29)* was re-analyzed to quantify expression of *Punc*, *Nope*, and *Prtg* in sensory and motor neuron subpopulations of interest. Using available data from three biological replicates at E12.5, read alignment, barcode and UMI counting, and aggregation was performed using the Cell Ranger pipeline (version 7.0.0, 10X Genomics), which uses STAR aligner *(51)*. The resulting barcode-UMI matrix was processed in R via the standard Seurat workflow (version 4.1.0) *(52)*. Cells which contained more than 6% UMI counts associated with mitochondrial genes or which expressed fewer than 500 genes were excluded from analysis, leaving a total of 13,021 cells that were analyzed. Cells were classified as neurons based on expression of *Tubb3* or *Elavl3*. Sensory neurons were further defined by expression of *Six1* or *Tlx2* and then stratified based on their expression of Trk receptors. Single-positive populations for each of the three Trk receptors were identified (193 *TrkA^+^*cells, 41 *TrkB^+^* cells, 84 *TrkC^+^* cells), as well as populations where these markers were expressed combinatorially (80 *TrkA^+^/TrkC^+^* cells, 46 *TrkB^+^/TrkC*^+^ cells, 72 *TrkA^+^/TrkB^+^/TrkC^+^* cells; because the *TrkA^+^/TrkB^+^* population contained fewer than 10 cells, it was excluded from analysis). Motor neurons were identified by *Mnx1* expression and further subdivided based on the brachial-level motor column markers *Foxp1* (LMC; 44 cells) and *Lhx3* (MMC; 16 cells). All expression-based classification of cell types used a UMI threshold of 1.

Expression levels of *Punc*, *Nope*, and *Prtg* were quantified for each subpopulation, and normalization was performed by dividing the transcript count for the given gene by the cell’s total transcript count.

### Expression constructs and recombinant proteins

Information about plasmids for protein expression is listed in Table S1. Untagged and AP-tagged proteins for the AP fusion protein binding assays and explant attraction/repulsion assays were expressed in COS-7 or HEK293 cells as previously described *(49)*. For use in the protein microarray screen, a fusion protein of the mouse [m]Punc ECD to the Fc portion of human IgG was generated at Genentech. For SPR experiments, we generated baculoviral transfer plasmids based on pAcGP67A. These plasmids were co-transfected with linearized baculoviral DNA (BestBac 2.0 (Expression Systems)) into Sf9 cells to produce baculoviruses using the Cellfectin II Reagent (Gibco). High Five cells were infected with the baculoviruses for expression, and proteins were purified from media using affinity chromatography with Ni-NTA Agarose resin (Thermo Scientific), and size-exclusion chromatography with a Superdex 200 Increase 10/300 column (GE Healthcare) in HBS (HEPES-buffered saline: 10 mM HEPES, pH 7.2 and 150 mM NaCl).

### Protein microarray screen

Details pertaining to the secreted protein microarray production, screening methodology, slide processing, and data analysis have previously been described *(32)* . For the protein interaction screen, mPunc-ECD-Fc, a fusion of mNope ECD (G22-H953) to human Fc (Biotechne R&D Systems 1394-NP), and a fusion of mPrtg ECD (F36-A952) to a His_6_ tag (Biotechne R&D Systems 6795-PR) were used as baits. To form protein A microbead-Fc fusion complexes for screening, a 1:1 mixture of the Fc fusion bait proteins and Cy5-labeled-hIgG was allowed to complex with multivalent protein A microbeads (Miltenyi Biotech). His-tagged Prtg protein was labeled directly with Cy5 monoreactive dye (GE Healthcare). Each bait protein was screened in duplicate against a total number of 1241 secreted proteins distributed across two microarray slides. Blocking, incubation, and wash steps were performed on an automated a-Hyb hybridization station (Miltenyi Biotech). Processed slides were scanned for Cy5 fluorescence using a GenePix 4000B scanner (Molecular Devices) and analyzed using GenePix Pro 6.0 software (Molecular Devices). Data analysis to identify hits was carried out as described before *(32)*.

### Cell culture and transfection

COS-7 and HEK293 cells were grown in on plastic dishes in high-glucose DMEM (Gibco) supplemented with 10% fetal bovine serum (Gibco) and 1x penicillin/streptomycin/glutamine (P/S/G) (Gibco). Cells were transfected in Opti-MEM (Gibco) using Viofectin (Viogene) according to the manufacturer’s recommendations.

### AP fusion protein binding assays

AP fusion protein binding assays were performed as previously described, with receptors being expressed directly in COS-7 cells and AP-tagged ligands being produced in HEK293 cells and harvested from conditioned OptiMEM media *(53)*.

### Surface plasmon resonance

For SPR experiments, mWFIKKN2 was N-terminally biotinylated using Sulfo-NHS-SS-Biotin (Thermo Scientific), where a 20x molar excess of dissolved biotin were mixed with purified mWFIKKN2 in MES-Buffered Saline (50 mM MES, pH 6.5, 150 mM NaCl) overnight. We used a Streptavidin (SA) Series S chip on a Biacore 8K (Cytiva) in HBS supplemented with 0.05% Tween-20 and 0.1% BSA. Five channels in the SA Series S chip were used, one for each analyte. Varying Response Units (RU) between ≈108-135 RUs of biotinylated mWFIKKN2 were captured on the SA chip channels. Titration series of the five receptors were injected on the chip with 60 seconds for the fast-binding Nope, and 150 seconds for Prtg, Punc, DCC and Neo1, followed by 300 seconds of dissociation for Nope, and 500 seconds for Prtg, Punc DCC, and Neo1. No regeneration of the chip surface was necessary for Nope, as this analyte fully dissociated, meanwhile injections of 2M MgCl_2_ were used for the rest of the analytes. The titration series included two pairs of duplicates, which showed no significant loss of chip activity through the titration series. All titrations and immobilization steps were performed at 25 °C. The steady-state responses were used to plot binding isotherms in Prism version 6. A binding model with no non-specific binding terms in Prism was used to calculate dissociation constants.

### Neuronal explant culture

Dorsal root ganglion (DRG) and ventral spinal cord explants from E12.5 and E11.5 mouse embryos, respectively, were dissected in L-15 media (Gibco) with 5% horse serum and cultured in collagen gel as previously described *(44)*. Aggregates of COS-7 cells were prepared the day after transfection and placed 300-500 μm from tissue explants. COS-7/DRG cocultures were grown for 18-24 hours in DRG media (Neurobasal, 1x P/S/G, 2% B27 (all Gibco), 0.5% glucose, 50 ng/ml NGF (Promega)). COS-7/ventral spinal cord cocultures were grown for 44-48 hours in motor neuron media (Neurobasal-A (Gibco), 2% B-27, 1x P/S/G, 0.5% glucose). To assess sensory axon fasciculation on a 2D substrate, DRG explants from E12.5 mouse embryos were dissected as described above and cultured for 18-24 hours in DRG growth medium on 8-chamber glass slides coated with 1 mg/ml poly-D-Lysine (PDL) (Sigma) and 5 μg/ml laminin (Millipore).

### Immunohistochemistry

All spinal cord sections were collected from brachial level. IHC on 150-μm-thick vibratome sections *(50)*, 2D neural tissue explants *(50)*, collagen gel COS-7 cell cultures *(49)*, and Dunn chamber cultures *(53)* were described previously, with blocking buffer used for IHC on Dunn chamber cultures and vibratome sections altered to include 2.5% donkey serum in lieu of bovine serum albumin. Primary antibodies used for IHC were goat polyclonal antibodies against TrkA, TrkB, and TrkC (R&D Systems, 1:200), rabbit polyclonal antibodies against TuJ1 (Biolegend, 1:500), Peripherin (Millipore, 1:200), and FoxP1 (Abcam, 1:500), mouse monoclonal antibodies against NF (DSHB, 1:200) and Islet1/2 (DHSB, 1:200), and chick polyclonal antibodies against GFP (Abcam, 1:200). Secondary antibodies were Alexa488-conjugated donkey anti-goat, Alexa647-conjugated donkey anti-goat, Alexa488-conjugated donkey anti-rabbit, Alexa647-conjugated donkey anti-rabbit, Alexa488-conjugated donkey anti-mouse, and Alexa594-conjugated donkey anti-mouse (all Invitrogen; 1:200), as well as Dylight405-conjugated donkey anti-mouse, DyLight405-conjugated donkey anti-rabbit, and Alexa647-conjugated donkey anti-chicken (all Jackson ImmunoResearch Laboratories, 1:200). Hoechst 33342 (Molecular Probes, 1:1000) was added with the secondary antibodies. Images were acquired on a Nikon Ti-E microscope or Nikon CSU-W1 SoRa Ti2-E microscope.

### Quantification of axon growth from explants in vitro

A minimum of 3 independent experiments were performed to quantify axonal growth from neuronal explants. For each replicate of a given experimental condition (n = 1), between 4 and 17 Tuj1/Peripherin-stained ventral horn explants or Tuj1/TrkA-stained DRG explants were used for quantification, and analysis was performed blinded to experimental condition. Axonal bundles were traced using the NeuronJ plugin for FIJI and measured from the point where they first emerge from the explant to their distal tip, and they were sorted into quadrants proximal, intermediate, and distal to COS-7 aggregates as described previously *(44, 49)*. For DRG explants, the total summed lengths of TrkA^+^ axons in each quadrant for all explants relative to COS-7 aggregates were determined and divided by the total summed length from all quadrants of all explants in each experiment. For ventral spinal cord explants, a similar analysis was performed using Peripherin as marker for motor axons. The means for relative proximal and distal growth from several independent experiments were compared in a paired two-tailed *t* test (n and *p* are indicated in figure legends). The mean total summed motor axon lengths from each condition across several independent experiments were also compared in a paired two-tailed *t* test (n and *p* are indicated in main text).

### Quantification of axonal fasciculation in vitro

Quantification of in vitro sensory axon fasciculation was performed as previously described for motor axons *(50)*. Between 8 and 12 explants per embryo were analyzed, with 3 embryos (n) per genotype. Analysis was performed blind to genotype. In short, the diameters of all Tuj1^+^ fascicles at a distance of 100 μm away from the edge of the explant were determined using the automated measurement tool in the NIS Elements software (Nikon), and correct image segmentation for measurements was confirmed by eye. Five diameter bins were defined (0-2 μm, 2-5 μm, 5-10 μm, 10-15 μm, and 15+ μm), and the total number of fascicles in each bin was calculated. These counts were then normalized to the total number of Tuj1^+^ fascicles for all explants for the entire experiment, so that the number of bundles within each bin was expressed as a percentage of total fascicle number per biological replicate. Normalized fascicle numbers for wild-type and *WFIKKN2* mutant animals for each bin were compared in an unpaired two-tailed *t* test (n and *p* are indicated in figure legends).

### Dunn chamber axon turning assay

Dunn chamber axon turning assays were adapted for sensory and motor neurons but essentially performed as previously described for spinal commissural neurons *(53)*, with a few modifications. DRGs and ventral spinal cord were dissected and dissociated as previously described at E11.5 and E10.5, respectively *(53)*. Cells were plated on nitric acid-washed and baked 18-mm coverslips coated with 100 μg/ml PDL and 5 μg/ml laminin, cultured in DRG or motor neuron media (see section on neuronal explant culture), and used for experiments 16-26 h after plating. The age of neurons at the time of the experiment was therefore E11.5 + 1 DIV (equivalent to E12.5) for DRGs and E10.5 + 1 DIV (equivalent to E11.5) for motor neurons. For generation of dose-response curves and Nope-ECD-Fc blocking experiments, neurons from multiple wild-type embryos were pooled for each experimental replicate. For experiments comparing *Nope* mutant and wild-type littermate controls, neurons from individual embryos were dissociated and cultured independently. Media used for pre-culturing of neurons were reused in Dunn chambers. Recombinant mWFIKKN2 (Biotechne/R&D Systems, 2070-GS) was applied (at indicated concentrations) to the Dunn chamber outer well, and ≈30-40 visual fields covering the bridge region of each chamber were imaged repeatedly over 2 h. For Nope-ECD-Fc blocking experiments, recombinant mNope-ECD-Fc protein (Biotechne/R&D Systems, 1294-NP) was added at 25 μg/ml (≈ 0.2 μM) to WFIKKN2-containing media (100 ng/ml ≈ 2 nM for sensory neurons, 1000 ng/ml ≈ 0.02 μM for motor neurons) immediately prior to Dunn Chamber assembly. Images were acquired on a Nikon Ti-E microscope.

### Quantification of axon turning in Dunn Chambers

Quantitative analysis of axon turning in Dunn chambers was performed as described previously *(53)*. All analyses were done blinded to genotype and experimental conditions. Neuronal identity was confirmed by post-hoc immunostaining of the imaged coverslip for the sensory axon marker TrkA in DRG cultures, or the motor neuron markers Islet1/2 and FoxP1 to differentiate between motor neuron subpopulations as previously described *(50)*. For each experimental replicate, axon turning angles from all analyzable neurons were averaged, and means across multiple replicates per condition or genotype were analyzed for statistical significance using either a one-way ANOVA with post-hoc Holm’s test for multiple comparisons (α = 0.05), an unpaired two-tailed *t-*test for *Nope* mutant versus wild-type comparisons, or a paired two-tailed *t-*test for other comparison between two conditions (n and *p* are indicated in figure legends).

### qRT-PCR

To determine *Nope* mRNA abundance, whole E12.5 *Nope* knockout embryos and wild-type littermates were homogenized at 4 °C in RNA STAT-60 (Tel-Test), and RNA was isolated according to the manufacturer’s recommendations. After treatment with DNase I, RNA was reverse transcribed using iScript cDNA synthesis kit (Bio-Rad). cDNA was used for quantitative PCR with SsoAdvanced Universal SYBR Green Supermix (Bio-Rad #1725270) on a StepOne Plus thermocycler (Applied Biosystems). The primers used were CATGGCCTTCCGTGTTCC and CAGTGGGCCCTCAGATGC for *GAPDH*, TGACATGGAGCAAGGATGGA and AGCCTGCATCACTGTCCTG for *Nope* exons 2 through 4, GGACAGCACGCTTTGAATG and ACAAAGTGTACCGAGTCCGG for *Nope* exons 8 through 10. The amplification efficiency of PCR reactions was determined by analyzing serial dilutions of samples, and each reaction was performed in duplicate. Relative *Nope* expression levels were normalized to *GAPDH* expression, and the means from 2 embryos of each genotype were determined.

### Quantification of ectopic axon projections at the motor-sensory plexus

TuJ1-stained vibratome sections of E12.5 embryo were imaged and maximum-intensity projections of z-stacks were generated for quantification. Sections were only analyzed if they captured the motor-sensory plexus (MSP) as well as the dorsal and ventral ramus. Analysis was performed blinded to genotype. The MSP was defined from the point where axons from the spinal cord and the peripherally-projecting root of the DRG converge, and a triangular region of interest (ROI) was generated by measuring 200 μm from the plexus following along both the dorsal and ventral ramus. Measurements of axon fascicles in this area were made from the point where axons emerged from the main nerve bundles and covered the entire arbor within the ROI. Using the FIJI plugin NeuronJ, the total summed length of all Tuj1^+^ axons within the ROI was quantified per hemisection. Values from 8-16 hemisections per embryo of each genotype (each embryo being a biological replicate, n = 1) were averaged, generating a single mean value per n. The averages across biological replicates within each experimental group were calculated and compared using unpaired two-tailed t tests (n and *p* are indicated in figure legend).

---

**Fig. S1.**
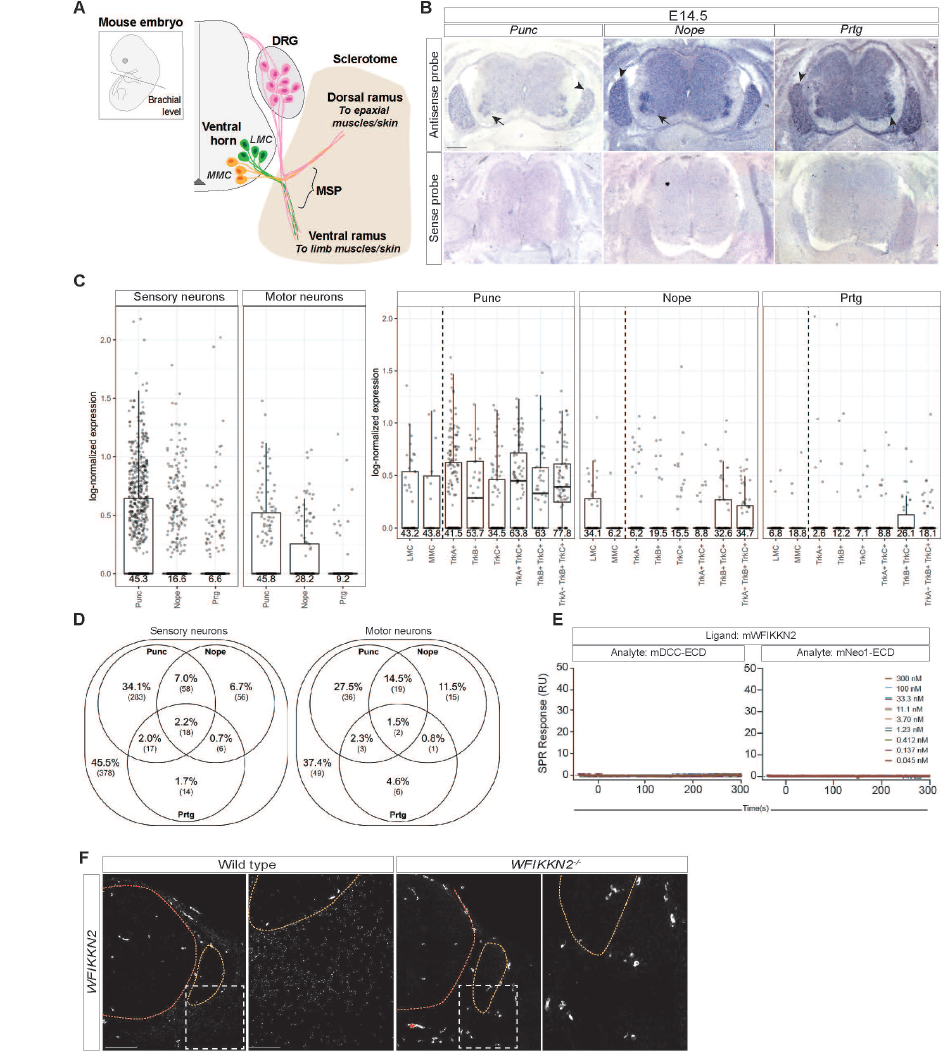
Binding and expression of WFIKKN2, Punc, Nope, and Prtg. **(A)** Schematic of transverse spinal cord hemisection at brachial level. Axons from motor neurons in the medial (MMC) and lateral (LMC) motor columns in the ventral horn and sensory neurons from the DRG converge at the MSP outside of the spinal cord before segregating into the dorsal and ventral ramus, which extend through the sclerotome within the developing somite. (**B**) Brachial spinal cord transverse sections from E14.5 mouse embryos were used for chromogenic in situ hybridization to detect *Punc, Nope,* and *Prtg* mRNA. All three genes show strong expression in DRGs (arrowheads) and the ventral horn (arrows); weak, non-specific signal is seen with sense probes. (**C**) Log-normalized expression of *Punc*, *Nope*, and *Prtg* transcripts at E12.5 in all sensory and motor neurons and specific sensory and motor neuron subpopulations, as extracted from published scRNA-Seq data *(29)*. Normalized transcript counts were multiplied by a scale factor of 10000 and then natural-log transformed using log1p. Box-and-whisker plots show the quartiles of each log-normalized expression distribution, with each point representing one cell. Numbers below each box represent the percentage of cells in that category that express at least one transcript of the given gene. Vertical dashed lines separate motor neuron subpopulations from sensory neuron subpopulations. (**D**) Venn diagrams showing coexpression of *Punc*, *Nope*, and *Prtg* transcripts at E12.5 in sensory and motor neurons in published scRNA-seq data *(29)*. For each partition, the percentage of the total population is shown, as well as the corresponding cell count. (**E**) WFIKKN2-DCC and WFIKKN2-Neo1 binding was studied by SPR. Shown are SPR sensograms for mDCC/Neo1-ECDs as analytes and mWFIKKN2 as ligand. Legend refers to concentration of analyte injected on the SPR chip. (**F**) Upper brachial spinal cord sections from E12.5 wild type and *WFIKKN2^−/−^* embryos were used for RNAScope fluorescent in situ hybridization to demonstrate specificity of *WFIKKN2* probe signal. No *WFIKKN2* mRNA is detected on *WFIKKN2* mutant sections. Spinal cord and DRG outlined with red and orange dotted lines, respectively. Zoomed-in regions are indicated by white dotted outlines. Asterisk indicates non-specific labeling from blood vessels. Scale bars, (B) 200 μm; (F) 100 μm; high magnification 50 μm. Error bars indicate SEM.

**Fig. S2.**
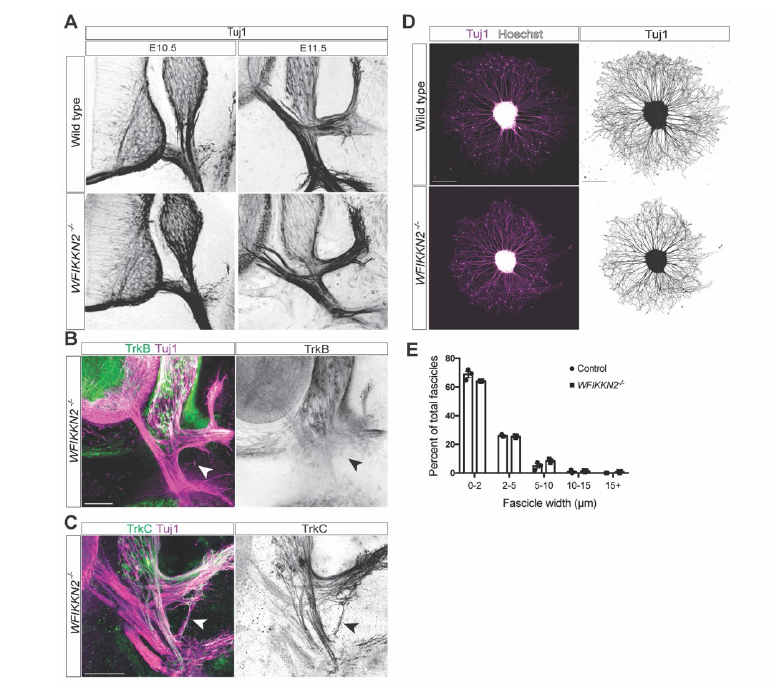
Sensory axon guidance defects in mice lacking WFIKKN2. **(A)** Brachial-level transverse sections of E10.5 and E11.5 wild-type and *WFIKKN2^−/−^* embryos were stained for Tuj1. Peripheral axon organization in *WFIKKN2^−/−^*animals is comparable to wild-type littermates. (**B** and **C**) Transverse sections of E12.5 *WFIKKN2^−/−^* embryos were labeled with Tuj1 antibody, Hoechst stain, and either TrkB antibody (B) or TrkC antibody (C). Aberrantly projecting axons (black arrowhead) are positive for TrkC, but not TrkB. (**D** and **E**) E12.5 DRG explants from wild-type and *WFIKKN2^−/−^* embryos were cultured on a 2D substrate and labeled with TuJ1 antibody and Hoechst stain (D), and thickness of axon bundles was quantified (E). Loss of WFIKKN2 does not affect axon fasciculation (n = 3 biological replicates, bin 0-2 μm *p* = 0.0964, 2-5 μm *p* = 0.46 5-10 μm, *p* = 0.1201, 10-15 μm *p* = 0.1808, >15 μm *p* = 0.1161). Scale bars (A-D) 100 μm. Error bars indicate SEM.

**Fig. S3.**
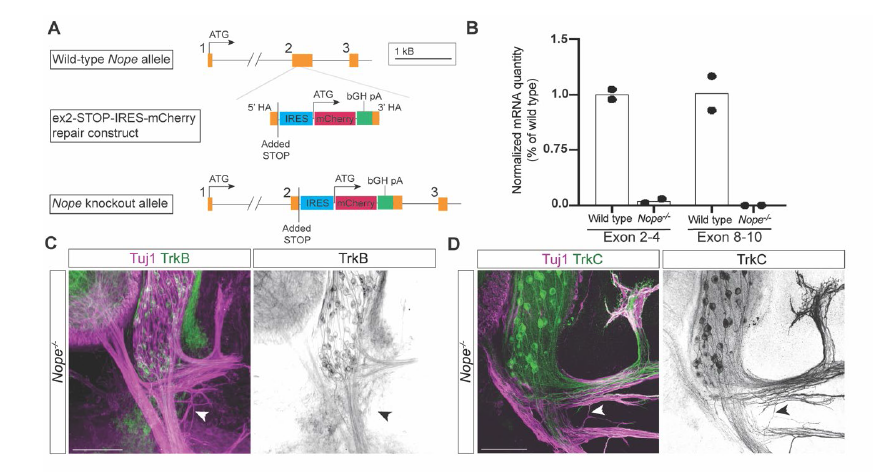
Generation and analysis of *Nope* knockout mice. **(A)** A schematic showing CRISPR/Cas9-mediated inactivation of *Nope* by targeted insertion of a STOP-IRES-mCherry-polyA cassette into exon 2 of the *Nope* gene. (**B**) *Nope* mRNA abundance in E12.5 *Nope^−/−^*embryos and wild-type littermates was determined by qRT-PCR for exons 2-4 and exons 8-10. *Nope* mRNA levels in the *Nope* mutant are reduced by >95% when compared to wild type (n = 2 embryos/genotype). (**C** and **D**) Brachial-level transverse sections of E12.5 *Nope^−/−^* embryos were labeled for TuJ1 and either TrkB (C) or TrkC (D). Aberrantly projecting axons (arrowhead) are positive for TrkC, but not TrkB. Scale bars (C, D) 100 μm.

**Fig. S4.**
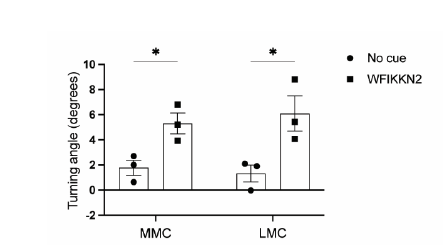
Motor axon attraction to WFIKKN2. Quantification of E10.5 + 1 DIV wild-type MMC and LMC motor axon turning angles in response to 1000 ng/ml WFIKKN2. WFIKKN2 attracts both MMC and LMC motor axons (n = 3 experimental replicates. MMC *p* = 0.0267, LMC *p =* 0.0378).

**Table S1.**
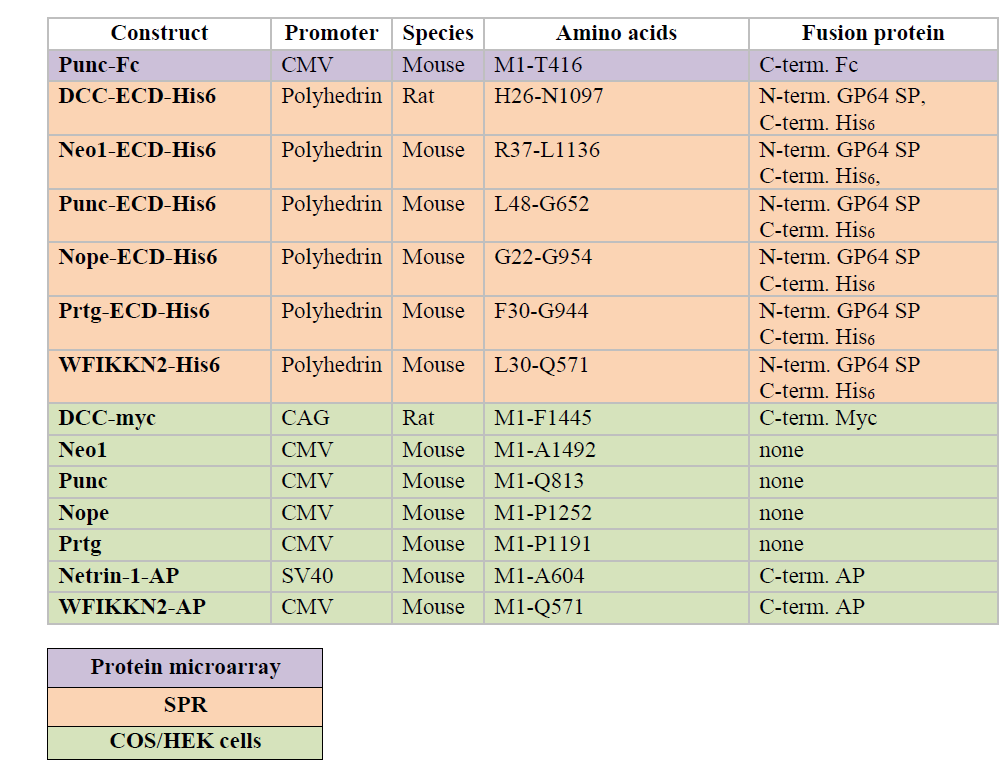
Protein expression constructs. Expression was driven with a Cytomegalovirus (CMV) promoter, a combined CMV/β-actin (CAG) enhancer/promoter, a Simian virus 40 (SV40) promoter, or a polyhedrin promoter. The species of origin and amino acid range for the cds of interest and the position of ectopic sequences and/or GP64 signal peptides (SP) fused to the cds are indicated. Legend indicates color coding of experimental use for each construct.

| **Construct** | **Promoter** | **Species** | **Amino acids** | **Fusion protein** |
| --- | --- | --- | --- | --- |
| **Punc-Fc** | CMV | Mouse | M1-T416 | C-term. Fc |
| **DCC-ECD-His6** | Polyhedrin | Rat | H26-N1097 | N-term. GP64 SP,<br>C-term. His<sub>6</sub> |
| **Neo1-ECD-His6** | Polyhedrin | Mouse | R37-L1136 | N-term. GP64 SP<br>C-term. His<sub>6</sub>, |
| **Punc-ECD-His6** | Polyhedrin | Mouse | L48-G652 | N-term. GP64 SP<br>C-term. His<sub>6</sub> |
| **Nope-ECD-His6** | Polyhedrin | Mouse | G22-G954 | N-term. GP64 SP<br>C-term. His<sub>6</sub> |
| **Prtg-ECD-His6** | Polyhedrin | Mouse | F30-G944 | N-term. GP64 SP<br>C-term. His<sub>6</sub> |
| **WFIKKN2-His6** | Polyhedrin | Mouse | L30-Q571 | N-term. GP64 SP<br>C-term. His<sub>6</sub> |
| **DCC-myc** | CAG | Rat | M1-F1445 | C-term. Myc |
| **Neo1** | CMV | Mouse | M1-A1492 | none |
| **Punc** | CMV | Mouse | M1-Q813 | none |
| **Nope** | CMV | Mouse | M1-P1252 | none |
| **Prtg** | CMV | Mouse | M1-P1191 | none |
| **Netrin-1-AP** | SV40 | Mouse | M1-A604 | C-term. AP |
| **WFIKKN2-AP** | CMV | Mouse | M1-Q571 | C-term. AP |

**Table S1. Protein expression constructs.**
|  |
| --- |
| **Protein microarray** |
| **SPR** |
| **COS/HEK cells** |

**Movie S1.** Two-hour time-lapse (30 frames/hour) of E11.5 + 1 DIV E12.5 DRG neuron cultured without cue in Dunn chamber. DRG sensory axons do not turn under control conditions. Scale bar, 25 μm.

**Movie S2**. Two-hour time-lapse of E11.5 + 1 DIV DRG neuron in a WFIKKN2 gradient (100 ng/ml) in Dunn chamber. DRG sensory axons turn away from of WFIKKN2. Scale bar, 25 μm.

**Movie S3.** Two-hour time-lapse (30 frames/hour) of E10.5 + 1 DIV motor neuron cultured without cue in Dunn chamber. Motor axons do not turn under control conditions. Scale bar, 25μm.

**Movie S4**. Two-hour time-lapse of E10.5 + 1 DIV motor neuron in a WFIKKN2 gradient (1000 ng/ml) in Dunn chamber. Motor axons turn toward WFIKKN2. Scale bar, 25 μm.

